# The Structure of the Drp1 Lattice on Membrane

**DOI:** 10.1101/2024.04.04.588123

**Authors:** Ruizhi Peng, Kristy Rochon, Anelise N. Hutson, Scott M. Stagg, Jason A. Mears

**Author notes:** R. Peng and K. Rochon contributed equally to this work. J.A. Mears and S.M. Stagg are joint senior authors.

## Abstract

Mitochondrial health relies on the membrane fission mediated by dynamin-related protein 1 (Drp1). Previous structural studies of Drp1 on remodeled membranes were hampered by heterogeneity, leaving a critical gap in the understanding of the mitochondrial fission mechanisms. Here we present a cryo-electron microscopy structure of full-length human Drp1 decorated on membrane tubules. Using the reconstruction of average subtracted tubular regions (RASTR) technique, we report that Drp1 forms a locally ordered lattice along the tubule without global helical symmetry. The filaments in the lattice are similar to dynamin rungs with conserved stalk interactions. Adjacent filaments are connected by GTPase domain interactions in a novel stacked conformation. We identified two states of the Drp1 lattice among the heterogenous dataset representing conformational changes around hinge 1. Additionally, we observed contact between Drp1 and membrane that can be assigned to the variable domain sequence. Together these structures revealed a putative mechanism by which Drp1 constricts mitochondria membranes in a stepwise, “ratchet” manner.

**SUMMARY:** This study provides new insights into the structure of Drp1 on lipid membranes. A locally ordered Drp1 lattice structure is solved and reveals intermolecular contacts and conformational rearrangements that suggest a mechanism for constriction of mitochondrial membranes.

## INTRODUCTION

Mitochondria constantly undergo fission and fusion to regulate their morphology, distribution, and size. This dynamic equilibrium is pivotal for numerous cellular processes, including apoptosis, cell cycle progression, damage isolation, and the overall maintenance of mitochondrial health (Wasiak, Zunino, and McBride 2007; Otera, Ishihara, and Mihara 2013; Nunnari and Suomalainen 2012). Drp1, an evolutionary conserved cytosolic protein, stands as the primary regulator of mitochondrial fission. Genetic deficiencies or mutations in Drp1 result in pronounced mitochondrial anomalies, which are associated with severe neurodevelopmental delay (Bauer et al. 2023; Robertson et al. 2023; Nolden et al. 2022). Conversely, enhancement of Drp1 activity is implicated in neurodegenerative disorders such as Parkinson’s, Alzheimer’s, and Huntington’s disease (Wang et al. 2011; Yan et al. 2015; Haun et al. 2013).

Mitochondria fission by Drp1 is a multistep process. Previous studies showed that mitochondria constriction involves the endoplasmic reticulum and actin, followed by recruitment of Drp1 by Mff and MiD49/MiD51 to the outer mitochondrial membrane (OMM) in mammals (Osellame et al. 2016). A recent study showed a novel peripheral fission of mitochondria where Drp1 is recruited by Fis1 to the relative peripheral position (Kleele et al., 2021). Despite the different modes, after recruitment to mitochondria OMM, Drp1 is reported to oligomerize through interactions with membrane lipids forming ring shape structures (Ingerman et al. 2005; Fröhlich et al. 2013). GTP hydrolysis promotes the constriction of the ring structure, providing the contractile force needed to mediate mitochondrial division (Francy et al. 2015; Koirala et al. 2013). While Drp1 plays a central role in mitochondria fission, the conformational changes required to drive the assembly and constriction of this contractile machinery are poorly understood.

Mechanistic understanding of Drp1 is largely inferred from dynamin superfamily proteins (DSPs). Crystallographic studies showed that Drp1 has similar domain arrangements when compared with dynamin, including a highly conserved GTPase domain (G domain), connected by a bundle-signaling element (BSE) to a stalk region that is composed of coiled-coil sequences formed by the middle domain and GTPase effector domain (GED). At the tip of stalk region, Drp1 has a ~100 amino acid loop termed the variable domain (VD), which was largely removed in the crystal structure. Dynamin has been reported to form a helical lattice along membrane tubules, and great efforts have been made to reconstruct dynamin remodeled membrane structures using helical symmetry (Kong et al. 2018; Chappie et al. 2011; Sundborger et al. 2014; Zhang and Hinshaw 2001). Recently, multiple high resolution cryo-electron microscopy (cryo-EM) structures of dynamin1 helices revealed the molecular mechanism of dynamin helix formation and how these structures channel the energy of GTP hydrolysis to constrict membranes through a compression of helical symmetry (Kong et al. 2018; Liu et al. 2021; Jimah et al. 2024). However, similar studies with mitochondrial fission DSPs show an expansion of helical lattices upon GTP hydrolysis (Mears et al. 2011). This suggests that the constriction mechanism may be different for Drp1, which is likely due to the larger magnitude of constriction for these oligomers that encircle intact mitochondria. When incubated with liposomes *in vitro*, Drp1 can form helical lattices on membrane with diameters ranging from 50 - 150 nm (Fröhlich et al. 2013; Kalia et al. 2018; Francy et al. 2017; Macdonald et al. 2014) making it suboptimal candidate for helical symmetry reconstruction. Another effort was made by solving the cryo-EM structure of a linear co-filament formed by Drp1 and MiD49, where Drp1 elongated via interfaces in the central stalk region similar to interactions observed in dynamin 1 helices (Kalia et al. 2018). However, the molecular mechanism of Drp1 forming circular rings on lipid membranes is still under investigation as commonly used helical symmetry reconstruction methods struggle to overcome the challenge of heterogeneity in Drp1 (Francy et al. 2017).

We previously developed a technique called reconstruction of average subtracted tubular regions (RASTR) to handle the challenges of heterogeneity in tubular samples (Randolph and Stagg 2020). Unlike helical symmetry reconstruction, which requires the protein lattice to be globally ordered, RASTR aims to break down the whole lattice into individual surfaces and capture the locally ordered sections. Instead of reconstructing whole tubular segments, individual tubular surfaces are upweighted and extracted. The resulting sub-particles are refined without imposing symmetry allowing for resolution of regions of the structure that deviate from helical symmetry. Here, we have applied RASTR on membrane tubules decorated with Drp1 that are highly heterogeneous. This resulted in a reconstruction with features that could be unambiguously assigned to Drp1 domains, which allowed us to build an atomic model of the Drp1 lattice (Peng 2023).

## RESULTS

### Drp1 is locally ordered on GalCer tubes

DSPs have been shown to remodel liposomes into tubules *in vitro* (Zhang and Hinshaw 2001; Sweitzer and Hinshaw 1998). Electron microscopy studies showed that dynamins form an ordered helical protein lattice on membrane tubules. Given the success of the iterative helical real space reconstruction approach (Egelman 2007), we and other researchers have successfully used helical symmetry reconstruction to resolve the structures of DSPs (Francy et al. 2017; Kalia et al. 2018; Kong et al. 2018; Sundborger et al. 2014; Alvarez et al. 2017). However, structure determination of Drp1-decorated membranes has been impeded by sample heterogeneity. It was demonstrated that upon incubation of Drp1 with liposomes, protein-decorated tubules were formed, but these tubules exhibited significant variability in their diameters, complicating the application of helical reconstructions (Fröhlich et al. 2013; Kalia et al. 2018; Francy et al. 2015; Macdonald et al. 2016). To mitigate this sample heterogeneity, we employed galactosyl ceramide lipids (GalCer) as a substrate (Wilson-Kubalek et al. 1998). GalCer lipids inherently form membrane nanotubes with a consistent diameter of approximately 20 nm (inner lumen), thereby constraining the diameter of Drp1-decorated membranes. We incubated full-length Drp1 isoform 3 with negatively charged GalCer tubes in the presence of GMPPCP. Upon incubation with GalCer tubes, well-defined protein decorations were observed (Fig. S1 A). However, layer lines with features able to be indexed were still not observed in the Fourier transforms of the tubules (Fig. S1, B and C). Consequently, we hypothesized that the lattice structures formed by Drp1 exhibit only local order, lacking long-range organizational coherence.

To validate this hypothesis and determine the size of the locally ordered area, we employed the RASTR technique (Randolph and Stagg 2020) on the cryo-EM dataset of Drp1-coated tubules. Particles were first processed using RASTR with four distinct spherical mask radii, all oriented on the middle of the tubule. As the mask diameter was reduced, a progressively smaller region of interest (ROI) was extracted from the original particles. These particle stacks were then independently aligned and classified in two dimensions (Fig. 1 A). The largest ROI mask produced a 2D average with diffuse continuous stripes of density like a barber pole going up the tubule axis. With a smaller 250 Å ROI mask, individual puncta were observed breaking up the continuous stripes of density in the middle of the tubules. With ROI masks smaller than 185 Å, more details were observed such as individual stalk regions and adjacent G-domains. These findings corroborated the notion that the Drp1 lattice was only locally ordered on GalCer tubes. This analysis revealed why the full tubule particles were suboptimal candidates for helical symmetry reconstruction; however, when processed via RASTR using a mask radius that aligned with the area of local order, they exhibited well-aligned features.

**Figure 1.**
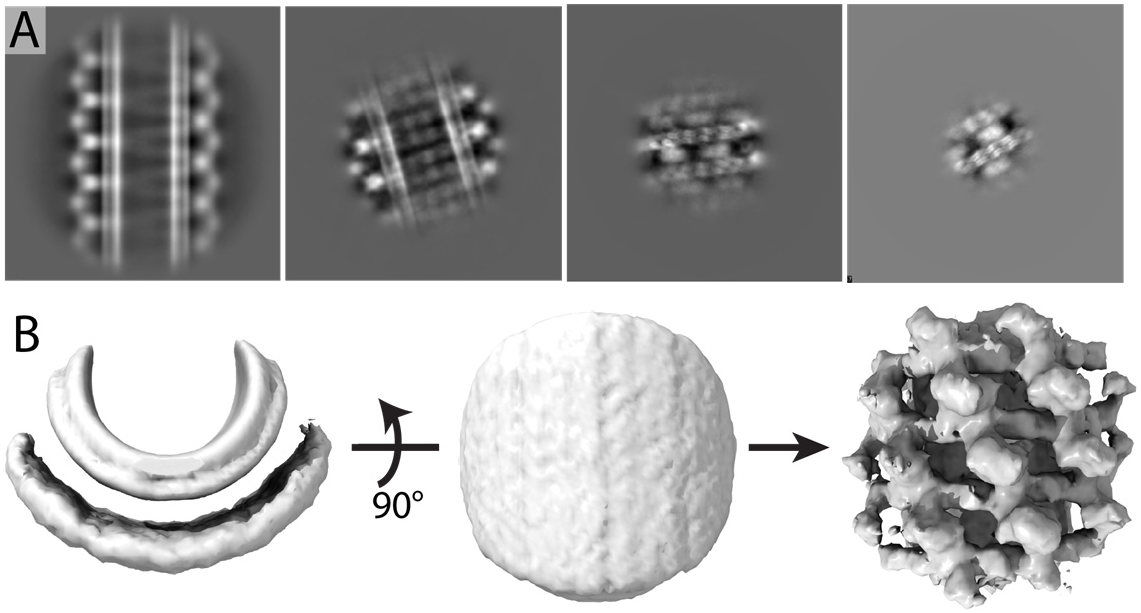
Drp1 lattice on GalCer tubes exhibit local order. **(A)** 2D classification of Drp1 particles with four different radii of ROI. From left to right, 425 Å, 250 Å, 185 Å, 130 Å. Box size, 888 Å. **(B)** Data processing for getting initial model. Left and middle, masked azimuthal average map. Right, Best 3D class map.

After selecting an appropriate ROI, we set about determining the 3D structure of the locally ordered Drp1 lattice. First, we needed an initial model for the single particle refinement without high-resolution features that might align spuriously to noise in the images, and the azimuthal average from the RASTR process provided a good initial model. Briefly, the azimuthal average was created by applying shifts and angle rotations to each particle image, making the tube axis at the center of the image and parallel to z axis. Images were rotated randomly around the tube axis, yielding the azimuthal average map (Fig. 1 B, left two panels). For Drp1, the azimuthal average map contained no high-resolution protein features but rather concentric layers of densities representing the two lipid bilayer leaflets and a smeared-out protein layer. After the azimuthal average initial model was made, we conducted 3D classification with eight classes using C1 symmetry in RELION. One class converged to a coherent structure, displaying distinct domain features (Fig. 1 B, far right density). These features, representing the G domain and stalk structures, exhibited repeats at a fixed angle twist and vertical rise, consistent with the canonical structural pattern observed in other DSPs (Kong et al. 2018; Liu et al. 2021). The fact that no symmetry was applied during refinement and that the initial model was featureless, providing no bias, demonstrated that the result was completely data driven. Based on this map, we estimated the helical symmetry parameters for the full helical filament. When we attempted a single particle reconstruction on full tubules with these parameters, the densities became smeared due to averaging of non-uniform features (Fig. S1 D), further demonstrating that Drp1 exhibits only local order when decorating GalCer tubes.

### Structure of the Drp1 Lattice on GalCer tubules

The resolution of the Drp1 map was improved with further local Euler angle refinement in *cis*TEM (Fig. S2 A). This resulted in a map at 15 Å resolution (Fig. 2 B, Fig. S2 B) that elucidated the features of the Drp1 lattice to a degree that enabled the construction of unambiguous atomic models (Fig. 2 C).

**Figure 2.**
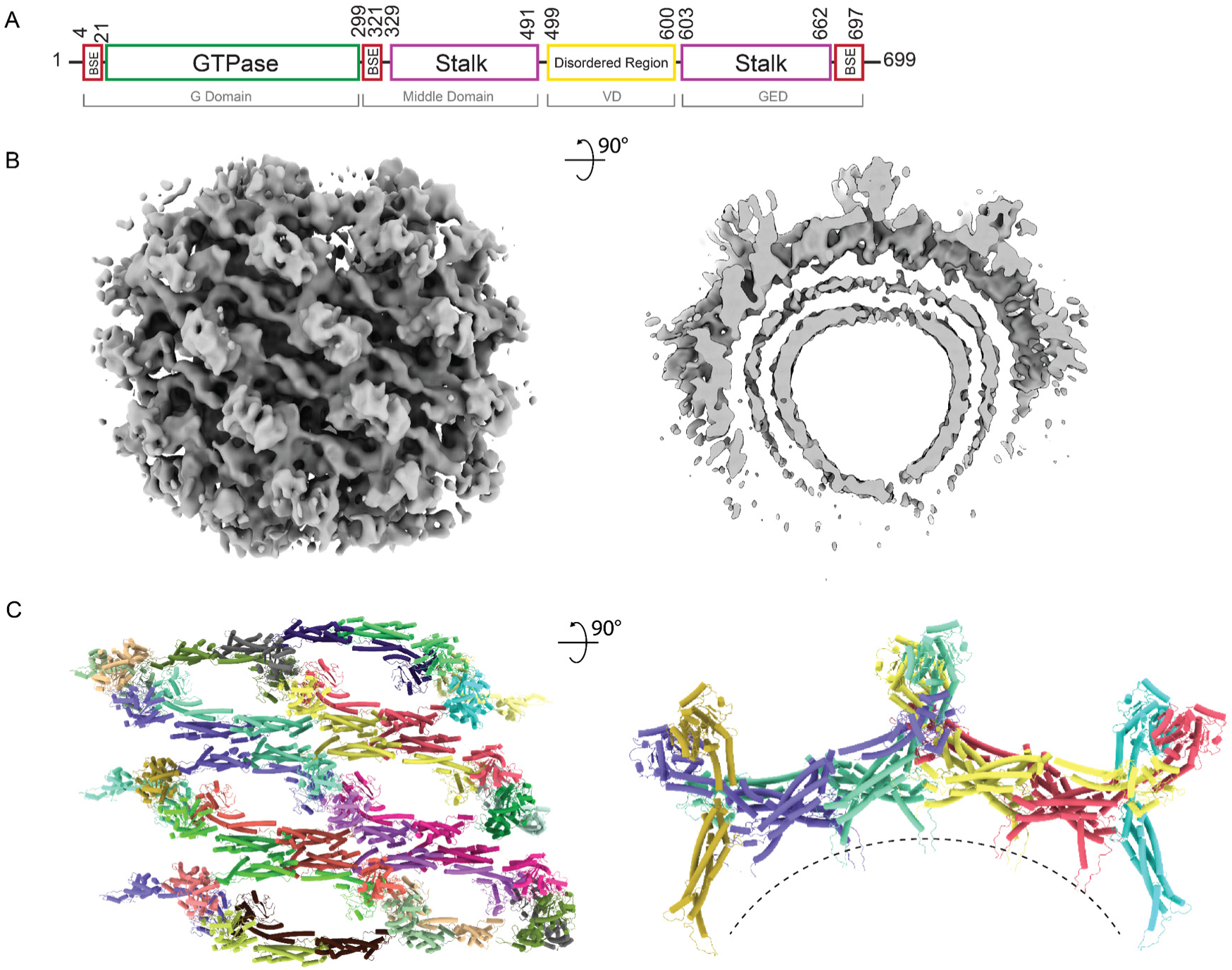
Architecture of assembled Drp1 in the GMPPCP-bound state on GalCer tubes. **(A)** Domain arrangement of Drp1. **(B)** Cryo-EM map of Drp1 decorated GalCer tubes. Image on the right is sliced to show one rung. **(C)** Lattice models. Image on the right is showing the center rung. Colored by individual dimers.

The Drp1 map showed features that were consistent with other DSPs. Specifically, a continuous filament was built via conserved intermolecular interfaces (numbered 1-3) between stalks (Reubold et al. 2015; Kalia et al. 2018; Kong et al. 2018; Liu et al. 2021). In our map, the best resolved region represented the stalk filament whose model could be fitted without ambiguity (Fig. S3 A). This elongated structure coordinated intermolecular contacts via the same interfaces (Interfaces 1-3, Fig. S3 D) previously reported (Kong et al. 2018; Kalia et al. 2018). In this way, the lattice builds through conserved interactions at well-defined interfaces.

### Conversely, differences in inter-domain organization were found in the BSE and G domain densities

For atomic modeling, we initially tried to fit solely the open form dimer (Kalia et al. 2018) (PDB: 5WP9) or the closed form dimer (PDB: 4BEJ). In both trials, half of the globular densities attributable to the G domains were unoccupied because one G domain in the dimer pair could not fit in those configurations. Therefore, we employed a strategy of fitting two states of dimers in the map. Using rigid body docking, the crystal structure (RCSB: 4BEJ) stalks aligned with minimal effort; however, it was clear the bottom G domains (Fig. 3, A and B) in the lattice would require a conformation where the BSE stretched further around hinge 1 than that observed in the crystal structure. The cryo-EM filament structure (RCSB: 5WP9) has an extended G domain conformation and was able to fit within the stalk density while still spanning the distance needed to reach the interface of the bottom G domains. A combination of two crystal structure monomers and two filament structure monomers were fit into the map by isolating the density representing the smallest asymmetric subunit of the lattice and removing the rest of the map, (Fig. 3 A). The interface aligning the four G domain monomers was positioned based on the structure of the dynamin 1 GTPase-GED (GG) fusion protein (RCSB: 3ZYC). This GMPPCP-bound dimer formed through G domain contacts that ensured a reasonable starting position comprised of conserved contacts. Once the asymmetric tetramer was fit to the best starting position, it was docked into the full map and additional monomers were added to build the stalk interfaces needed to constrain each monomer within the density. After minimization, Molecular Dynamics Flexible Fit (MDFF) was used to further refine the final asymmetric subunit.

**Figure 3.**
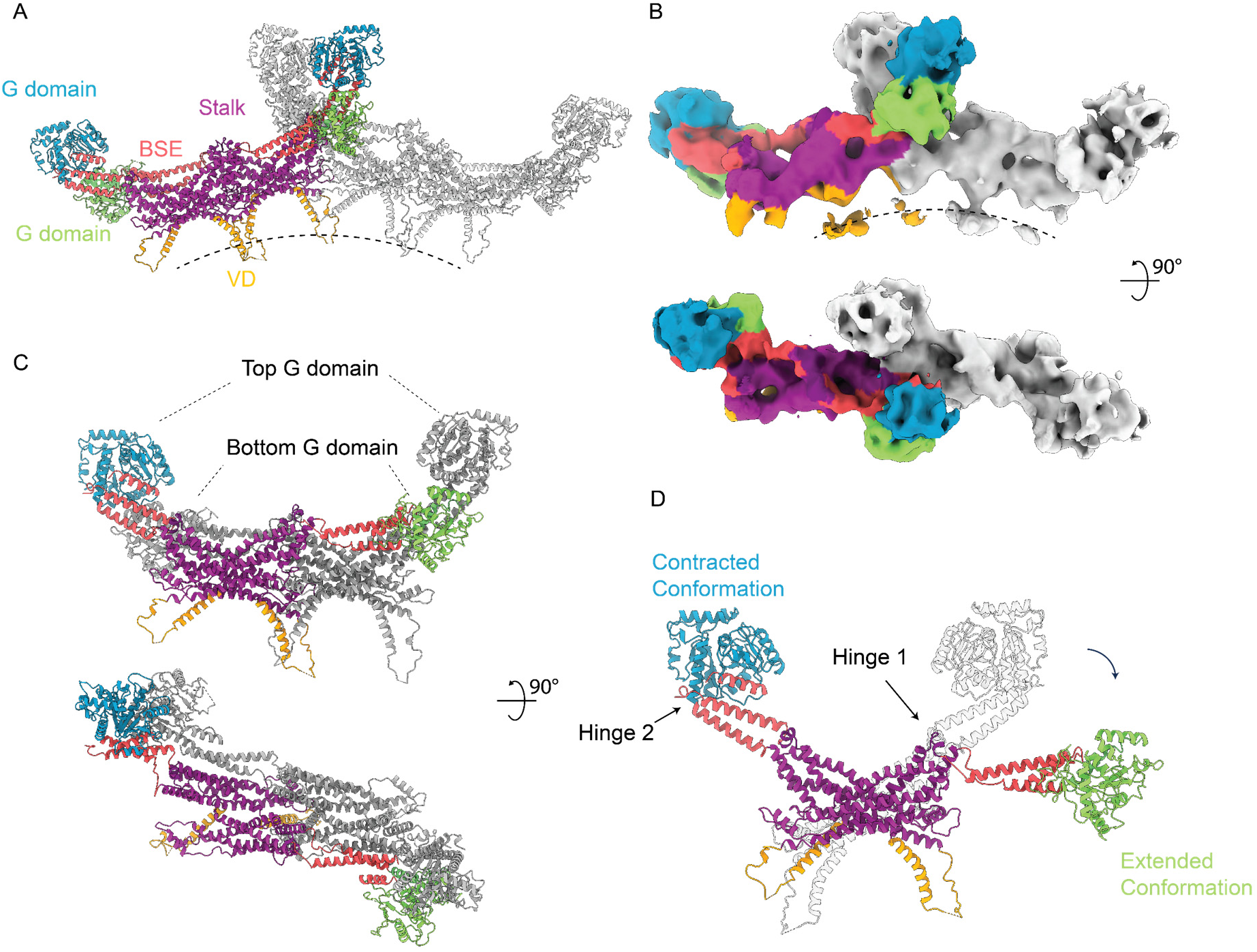
Asymmetric tetramer. Segmented models **(A)** and cryo-EM maps **(B)** showing two tetramers within same rung. One colored, another gray. **(C)** Asymmetric tetramer. Yellow and cyan representing two dimers. **(D)** Conformational change around hinge 1. Two dimers superimposed on stalk. Purple, stalk domain; Red, BSE; Green, bottom G domain in extended conformation; Blue, top G domain in contracted conformation; Brown, VD domain.

Surprisingly, the model converged to a configuration featuring G domains from two dimers stacked atop one another (Supplementary Video 1, Fig. 3 C, Fig. S3 B and C). Previous studies typically showed Drp1 dimers to be symmetric with both monomers exhibiting the same conformation around hinge 1 (Fröhlich et al. 2013; Kalia et al. 2018). However, in our model, each dimer possessed hinge 1 orientations in two distinct states, with one monomer in an extended conformation and the partner monomer in a contracted conformation (Fig. 3, C and D). This was consistent with a recent solution structure that showed that the hinge 1 loop in Drp1 is flexible, so the relative positions of the BSE and stalks are not restricted (Rochon et al. 2024). The extended conformation is consistent with the Drp1 filamet structure determined by cryo-EM (Fig. S4 B). Within the context of this tetrameric model, the interfaces between stalks remained consistent (Fig. S3 D). Another difference was the interface between G domains. In our model, the GG interface between the bottom G domains is the same as dynamin 1 (Fig. S3 E, green). The top GG interface is similar, but there is a ~10 Å shift (Fig. S3 E, blue) compared to the bottom interfaces or GG pairs reported in dynamin helices. This novel GG interface is the conformation that is most consistent with the observed density but is not definitive at the current resolution.

Altogether, the Drp1 lattice structure was characterized by parallel rungs. Each rung was elongated through oligomerization of the stalk domains in the same manner as reported before (Kalia et al., 2018). Interposed between these rungs, G domains served as contact sites that stabilized adjacent rungs (Fig. 2 C). The smallest repeating subunit was a tetramer, with two dimers stacking atop one another (termed the “stacked state”). The individual dimer in our structures was asymmetric, with one monomer in extended conformation containing a top position G domain and the other monomer in a contracted conformation containing a bottom position G domain.

### Conformational changes in Drp1 lattice

We collected another dataset in order to increase the resolution of the Drp1 lattice. However, while the stacked state lattice was identified in the new dataset, there was an increase in the heterogeneity, and the majority of particles shifted to an alternative state that we termed the “relaxed state” (Fig. 4 A, right). At the vertical center of this map, densities corresponding to stalks were observed, but the stacked G domain densities were absent in the relaxed state map (Fig. 4, A and B). Instead, a diffuse cloud of densities appeared at the radial position corresponding to the bottom G domains, while densities at the radial position of the top (i.e. most peripheral) G domains were not observed (Fig. 4 B). In the relaxed state of the Drp1 lattice, the central rung corresponding to stalk densities was consistent with the stacked state map, but stalk densities in adjacent rungs were less well-defined (Fig. 4 A). Additionally, we noted a curvature change between these two states. In the stacked state, the lipid membrane curvature exhibited local flattening at the center of the target region, which is similar to the larger diameter of Drp1 remodeled liposomes (Fig. S5 C), but in the relaxed state, the curvature at the center was higher (Fig. 4 C). We didn’t attempt to build a complete model of this map because of the diffuse and incoherent densities observed for the G domain cloud and adjacent rungs. However, in the center rung densities, it was clear that the stalk domain organization and interfaces were conserved between the two structures, indicating the stable intermolecular interactions between stalks. Hinge 1 in the relaxed state largely adopted the extended conformation and was not well-defined either due to heterogeneity or changes to the BSE extension.

**Figure 4.**
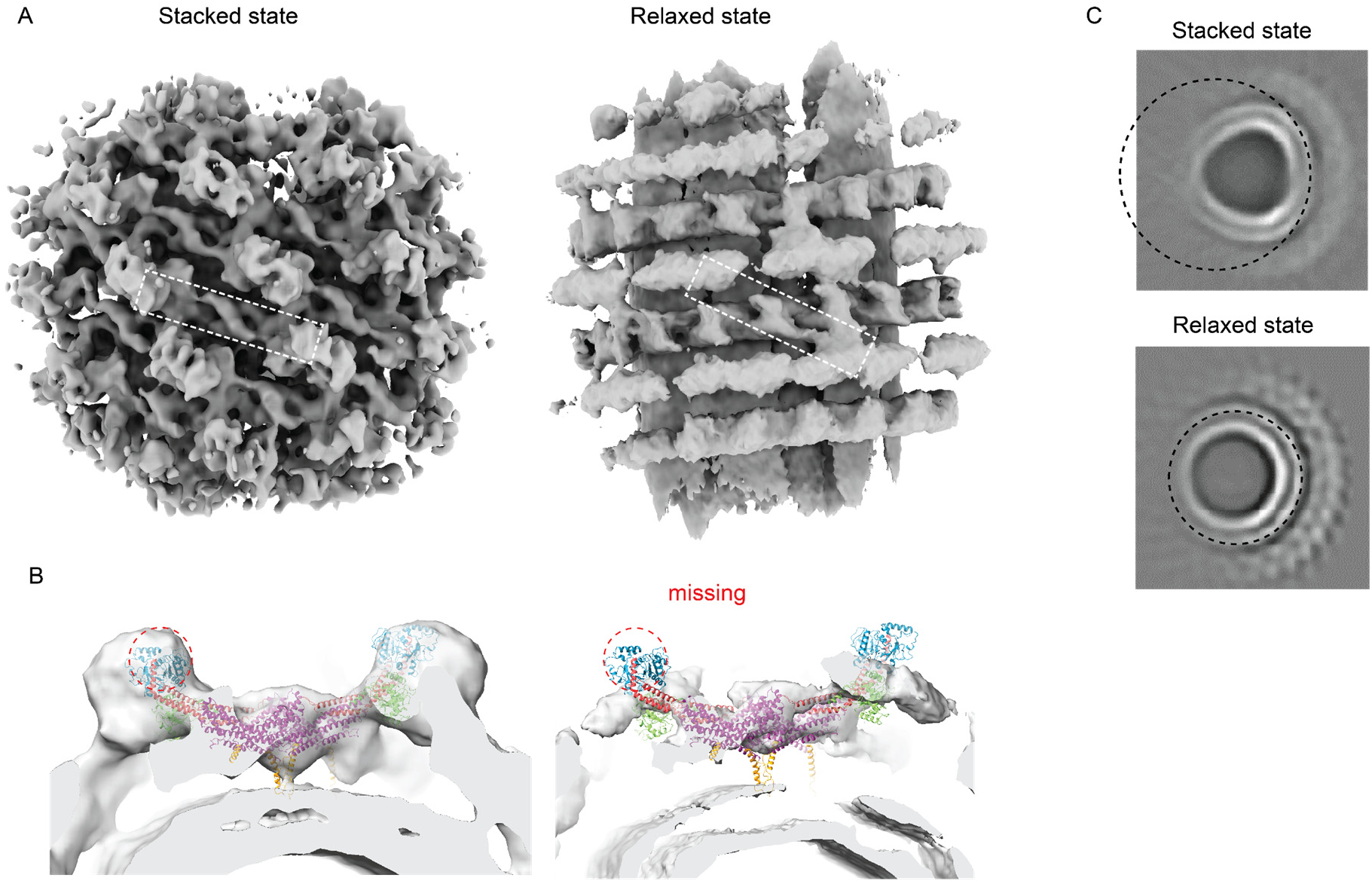
Comparison of Drp1 lattice in two states. **(A)** Cryo-EM map of Drp1 lattice. **(B)** Focused view showing one tetramer. Density of top G domain is missing in extended state map. **(C)** Projection of Cryo-EM map along tube axis. Local flattening of membrane densities in stacked state map. Left showing stacked state, right showing extended state. Purple, stalk; Red, BSE; Green, bottom G domain; Blue, top G domain; Brown, VD domain.

We further investigated the heterogeneity of the dataset using cryoDRGN, a machine learning based algorithm for characterization of heterogeneous cryo-EM structures (Zhong et al. 2021), and this revealed continuous flexibility in the Drp1 lattice. Analysis of the PC1 axis of the cryoDRGN latent space revealed a continuous shift from the stacked state lattice to the relaxed state lattice (Fig. S6 A, Supplementary Video 2). There is a strong correlation between the densities of the top G domains and the spacing of the rungs (Supplementary Video 2). When the top G domain density reached its highest occupancy, the spacing between Drp1 rungs was at its shortest, consistent with the stacked state lattice which contains well-defined G domain pairing. Conversely, at its lowest occupancy, the rung spacing expanded, indicative of the relaxed state lattice. Furthermore, when the top G domain was present, adjacent rungs exhibited features with similar resolution. However, in its absence, only the central rung had recognizable stalk features, while adjacent rungs appeared less ordered. Collectively, these findings suggest that the binding of G domains, especially with both top and bottom G domains, served to stabilize the entire lattice, maintaining the integrity of adjacent rungs. Conversely, compromised G domain binding weakened the connections between adjacent Drp1 lattice rungs, leaving the lattice flexible. The heterogeneity was further analyzed by cryo-electron tomography. While features representing stalk and bottom G domains were evident, the spacing between stalk features varied considerably (Fig. S7, A and B). Additional densities, corresponding to top G domains were occasionally observed extending from the bottom G domain (Fig. S7 C), but they were infrequent which is consistent with the cryoDRGN results.

The second cryo-EM dataset had a slightly different, though still saturating, GMPPCP concentration (2 mM compared to 1 mM originally), so we prepared cryo-EM grids with varying GMPPCP to see if that influenced the equilibrium between the stacked and unstacked states. Incubation with 0.5, 1, and 2 mM GMPPCP resulted in no change in the equilibrium. Therefore, we hypothesized that variations in the lipid reconstitution are what led to the shift in the equilibrium between our two datasets. Since the stacked state lattice appeared to flatten the membrane and PA is associated with increased membrane deformability (Zhou et al. 2024), we hypothesized that preparing membrane tubules with only PA and GalCer would shift the stacked/relaxed equilibrium. We prepared Drp1 coated tubules with 30% PA and no phosphatidyl ethanolamine (PE), and this shifted the equilibrium with an increase in the stacked state to 60.3% from 25.7% with PE and PA (Fig. S6 D). Thus, lipid composition appears to influence the stacked/unstacked equilibrium.

### Drp1 lattice contacts the lipid membrane

A novel feature in our stacked state cryo-EM map was the contact between the Drp1 protein lattice and lipid membranes (Fig. 5 A). The contact sites were distributed periodically on the lattice (Fig. 5 A), with each membrane contact coinciding with an individual tetramer. Further inspection revealed that these contact sites are located exactly under the proximal tips of the stalk at the center of tetramers, where the VD sequence starts (Fig. 5 A blue, Fig. 5 B top). We also observed extended densities in the region of the stalk tips between tetramers (VD at junctions between tetramers instead of the ones at the center of the tetramers), but these were of much lower occupancy or contact with lipid membrane densities was not observed (Fig. 5 A). Tracing back from the VD to monomers, we discovered that the VD structures at the centers of tetramers all originated from monomers with contracted conformations, whereas weaker VD interactions at intervening junctions all originated from monomers with extended conformations (Fig. 5 B). The different occupancies of the contact densities indicated a strong relationship between VD-lipid interactions and G domain conformation (Fig. 5 B). In support of this, we didn’t observe membrane contact in the relaxed state map (Fig. 4 B), where no G domains were in a contracted conformation. Together, the membrane interactions correlated with hinge 1 conformations in a pattern where the contracted conformation favored membrane connections and the extended conformation weakened this connection, which would permit lattice rearrangements (i.e. constriction).

**Figure 5.**
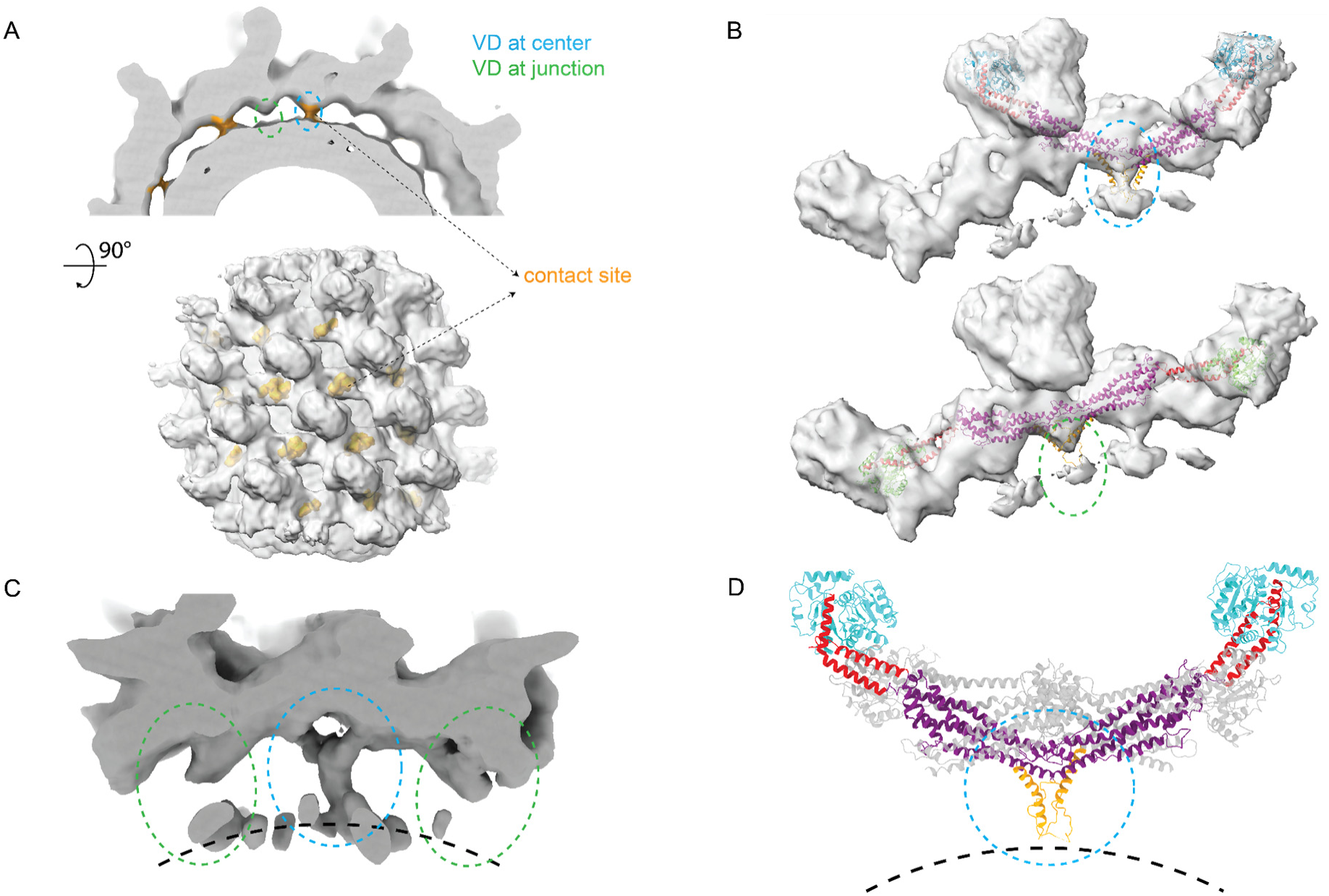
Contact between Drp1 lattice and lipids membrane. **(A)** Low-pass filtered cryo-EM map slice showing contact between Drp1 lattice density with lipids membrane density. Contact density colored by brown. Green dash circle indicating VD position of bottom G domain containing Drp1 monomer. Cyan dash circle indicate VD position of top G domain containing Drp1 monomer. **(B)** Focused view showing differences in membrane contact densities. Map are segmented from Fig. 2 A. Left, membrane contact within tetramers. Right, membrane contact between tetramers. **(C)** Cryo-EM map of sub-particle reconstruction at contact position. **(D)** Focused view of VD domain model. Monomer with top G domain conformation is colored. Monomer with bottom G domain conformation is colored gray.

The Drp1 VD is ~102 amino acids long and has been attributed to have a regulatory function by interacting with lipids (Adachi, Iijima, and Sesaki 2018; Adachi et al. 2016; Francy et al. 2015; Ugarte-Uribe et al. 2014). Several important charged residues in this region are associated with specificity for negatively charged lipid headgroups (Mahajan et al. 2021). However, the VD was either truncated or not well resolved in previous structural studies (Kalia et al. 2018; Fröhlich et al. 2013). Because the protein lattice interacts in a regular pattern in the stacked state map, we reasoned that the density for the VD may be apparent at lower resolution when compared to the rest of the map. Therefore, a sub-particle reconstruction strategy was employed to refine the VD proximal to the membrane, specifically focusing on the region at the center of a tetramer. In the resulting reconstruction, we observed densities extending from the terminal of the two paired stalks (Fig. 5 C). These densities extended towards membrane and fused together forming a thicker rod-like density (Supplementary Video 3). Notably, despite not enforcing symmetry during the refinement process, the resulting map showed a C2-symmetric density that was consistent with a pair of helices contributed by adjacent VDs.

The N-terminal end of the VD, which has been termed the Molecular Recognition Feature 1 (MoRF-1)(Mahajan et al. 2021) has been predicted to be helical. This helix at residues 499-520 was also predicted by AlphaFold (Jumper et al. 2021). MoRFs were found in intrinsically disordered regions (IDRs) and were capable of transitioning from a disordered to ordered under favorable conditions, like partner protein binding (Bugge et al. 2020; Fung, Birol, and Rhoades 2018). We tested the influence of the MoRF-1 region on the Drp1 lattice. Deletion of MoRF-1 sequence shifted the oligomer states of Drp1 in solution from a mix of dimer, tetramer, hexamer to mainly monomer and dimer (Fig. S8 B). Without MoRF-1, Drp1 also showed a highly impaired stimulation of GTPase activity when lipid was added (Fig. S8 A). Even with the deletion of MoRF-1, Drp1 readily associated with the membrane (Fig. S8 C), though the assembly appeared less ordered based on negative-stain EM images (Fig. S8 D). Therefore, we attributed the lack of stimulated GTPase activity on the destabilization of the lattice in the absence of MoRF-1. There were no published structures of Drp1 with a resolved density for VD. Based on the sub-particle reconstruction map, we docked a helix for the MoRF-1 into this contact position, forming an interface between the two adjacent helices in neighboring Drp1 monomers (Fig. 5 D). These coupled helices formed a local dimeric conformation that reached down to touch the surface of the lipid bilayer (Fig. 5 D).

## DISCUSSION

Here, we reconstructed the structure of the Drp1 lattice on membrane tubules. Despite the inherent flexibility of Drp1 particles, which renders them unsuitable for helical symmetric reconstruction, we demonstrated that localized order exists within the particles that can be extracted using RASTR and subsequently reconstructed. This approach revealed novel features about membrane bound Drp1 structure, including the discovery of stacked and relaxed states of the lattice, the development of a molecular model featuring an asymmetric tetramer as the repeating subunit, and the observation of membrane contacts in the VD that occur in the stacked state.

Previous studies on analogous membrane remodeling proteins often employed helical symmetry reconstruction (Zhang and Hinshaw 2001; Kalia et al. 2018; Kong et al. 2018; Nguyen et al. 2020), which requires long-range order in protein lattices. However, many membrane remodeling proteins are only locally ordered, reflecting the heterogeneous and flexible nature of membranes (Kong et al. 2018; von der Malsburg et al. 2023). Consequently, conventional cryo-EM reconstructions lack high-resolution features and/or have ambiguous helical order. Our current study demonstrates how RASTR addresses these challenges by isolating structural features from the ordered surfaces of protein lattices. Here, RASTR enabled the extraction of structural features from Drp1 that was not possible using conventional helical processing. The Drp1 lattice determined by RASTR exhibited parallel rungs indicative of the stalk densities, complemented by globular densities that correspond to the GTPase domains.

Building on these insights, we developed pseudoatomic models to elucidate the structural mechanisms underlying Drp1 assembly and membrane constriction. The resolved structure of Drp1 decorated membrane tubules revealed a novel stacked conformation formed by asymmetric tetramers. Each tetramer consists of two dimers, with two of the proteins in a contracted conformation and the other two in an extended conformation. These conformations differ primarily in the BSE and hinge 1 regions. In the extended state lattice, the BSE hinge swings outward, similar to the conformation observed in the filament structure of Drp1 paired with MiD49 (Kalia et al. 2018) and a truncated dynamin nucleotide-bound structure (Chappie et al. 2011) (Fig. S4 B). This structure also highlights the ability of hinge 1 to confer large conformational rearrangements of the G domain while still maintaining ordered stalk interfaces.

The stacked state for the G domains has not been observed previously in Drp1 reconstructions in the absence of membrane, nor has a similar state been observed in other DSPs. There are many pieces of evidence that support these novel results. First, the Drp1 lattice was resolved in the reconstruction after starting with the featureless azimuthal average as the initial model for refinement. No symmetry was imposed during refinement yet nearly identical features at periodic positions were resolved for the different asymmetric units in the resulting lattice. In this way, the map is self-validating. Another strong supporting piece of data is that the stacked state was recapitulated when classifying a second dataset that exhibited a greater degree of heterogeneity. Analysis of the heterogeneity revealed that the Drp1 lattice undergoes a breathing motion with continuous flexibility from a relaxed state where only one set of G domains in the Drp1 asymmetric tetramer are well-ordered to a compacted stacked state, where both sets of G domains are ordered and are stacked on top of one another. Together, these G domain interactions in the Drp1 lattice are more dynamic when compared to dynamin structures. In the dynamin structure (Kong et al. 2018), the G domain pairing is continuous and uniform. In the Drp1 lattice, the variation in inter-rung distance coincides with the altered pairing between adjacent G domains. In the relaxed state, only one of the G domain pairs is observed, suggesting that the other domains are unpaired, or irregular. This would be consistent with the weaker interactions between rungs that lead to variability in spacing. This G domain pairing is critical for the stimulated activity of DSPs on lipid membranes, and consistent with previous biochemical findings; the stimulation of Drp1 GTPase activity is much lower than that observed with dynamin (~10-15-fold stimulation for Drp1 compared to ~100-fold stimulation for dynamin) (Bustillo-Zabalbeitia et al. 2014; Macdonald et al. 2014; Warnock, Hinshaw, and Schmid 1996; Chappie et al. 2010).

We performed several experiments to characterize what factors influence the equilibrium between the stacked and relaxed states. We found that increasing the nucleotide concentration did not influence the stacked/relaxed equilibrium, but the equilibrium can be shifted towards the stacked state by varying lipid composition. When we increased or decreased GalCer% while maintaining the same PA/PE ratio, there were no significant differences, but when PE was removed the equilibrium shifted from 25.7% to 60.3% of particles in the stacked state. These results are consistent with our discovery of a membrane contact that was evident in the stacked state map but not apparent in the relaxed state map. Furthermore, our experiments removing the MoRF-1 region that formed the putative membrane contact in our map showed a significant alteration of Drp1’s behavior on lipid membranes. Together, these results suggest that lipids have an important role in regulating Drp1 lattice conformations, and the VD plays a role in sensing the lipid and transmitting the nature of this interaction within the lattice.

The transition between the two states appears to be mediated by G domain-G domain interactions. In the relaxed state map, it’s evident that adjacent rung densities are flexible as demonstrated by the diffuse densities in the map and by the continuous flexibility shown with cryoDRGN. Conversely, in the stacked state lattice, overall contacts between G domains are stronger, ensuring adjacent rungs are held firmly and closely. Consistent with this, adjacent rungs in the stacked state lattice are held by asymmetric GG contacts, but the top pair is lost in the relaxed state. Since this top G domain pair is in a contracted conformation, the natural changes that would follow disruption of this pair is extension of the G domain. This directionality is consistent throughout the lattice, and transition between the contracted and extended conformations would drive constriction of the Drp1 lattice as adjacent rungs ratchet through alternating G domain pairings (Supplemental Video 4). Consequently, throughout Drp1 tubules, some regions of the lattice will exhibit a relaxed state, while others will be stacked (Fig. S7, A and C).

Our RASTR reconstruction of the Drp1 stacked state lattice revealed that the initial segment of the VD becomes ordered and contacts the membrane. We discerned that in the stacked state lattice, the first ~20 residues in the VD occupy a vertical position, extending from the distal end of stalk to the outer leaflet of the membrane. Moreover, we discovered that the VD situated at the center of a tetramer exhibits a pronounced connection with the membrane, whereas the VD located at the junction between adjacent tetramers is sparsely populated (Fig. 5 A). This is reminiscent of the asymmetry observed in the dynamin lattice, where the PH domains impinge on the membrane depending on the conformation of the rest of the monomer. A similar phenomenon may be at play in the Drp1 lattice where the state of the G domain interactions is transmitted to the lipid-binding segment of the protein. In the same way that constriction of the lattice requires dynamic G domain interactions, the VD interactions with the membrane are likely dynamic in a coordinated manner that permits sliding on the membrane surface while still maintaining association.

Previous investigations into Drp1 proteins recognized the fission process as a continuous, dynamic process, with the diameter of Drp1 rings gradually decreasing. In the two states that we captured, two distinct curvatures demonstrated the inherent flexibility and dynamic nature of the lattice, potentially representing two intermediate states in the dynamic process. We posit that the cyclical transition of G domain pairing and VD-membrane interactions in transition from the extended state to the relaxed state in Drp1 underpins the sliding and contraction of the Drp1 lattice during mitochondrial remodeling (Fig. 6, Supplementary Video 4). In this model, upon recruitment to the mitochondria, Drp1 proteins extend through interactions at stalk interfaces. As the filament lengthens and encircles the membrane, adjacent rungs connect through the G domains, resulting in a stabilized structure that favors a flatter geometry, which may also explain the preference towards larger tubules with Drp1 when compared with dynamin (100+ nm diameter for Drp1 vs 50 nm for dynamin). The binding of GTP instigates a perpetual transition between the constricted and extended G domain states that facilitate breathing in the lattice from stacked to relaxed states. This conformational flux within the G domain propels the stalk-mediated filaments to slide in the direction of constriction that is biased by the right-handed nature of the lattice and the natural extension of the contracted G domain in the asymmetric dimer that will reach to form a new pair that promotes sliding between rungs and drive the constriction of the lattice. As the lattice approaches higher curvature, the strain on maintaining G domain connections will be greater. In agreement, Drp1 filaments dissociate from the membrane after constriction. It is possible that additional mitochondrial fission factors, including partner proteins and cytoskeletal elements, could stabilize the lattice at higher curvatures to enhance constriction and drive membrane fission.

**Figure 6.**
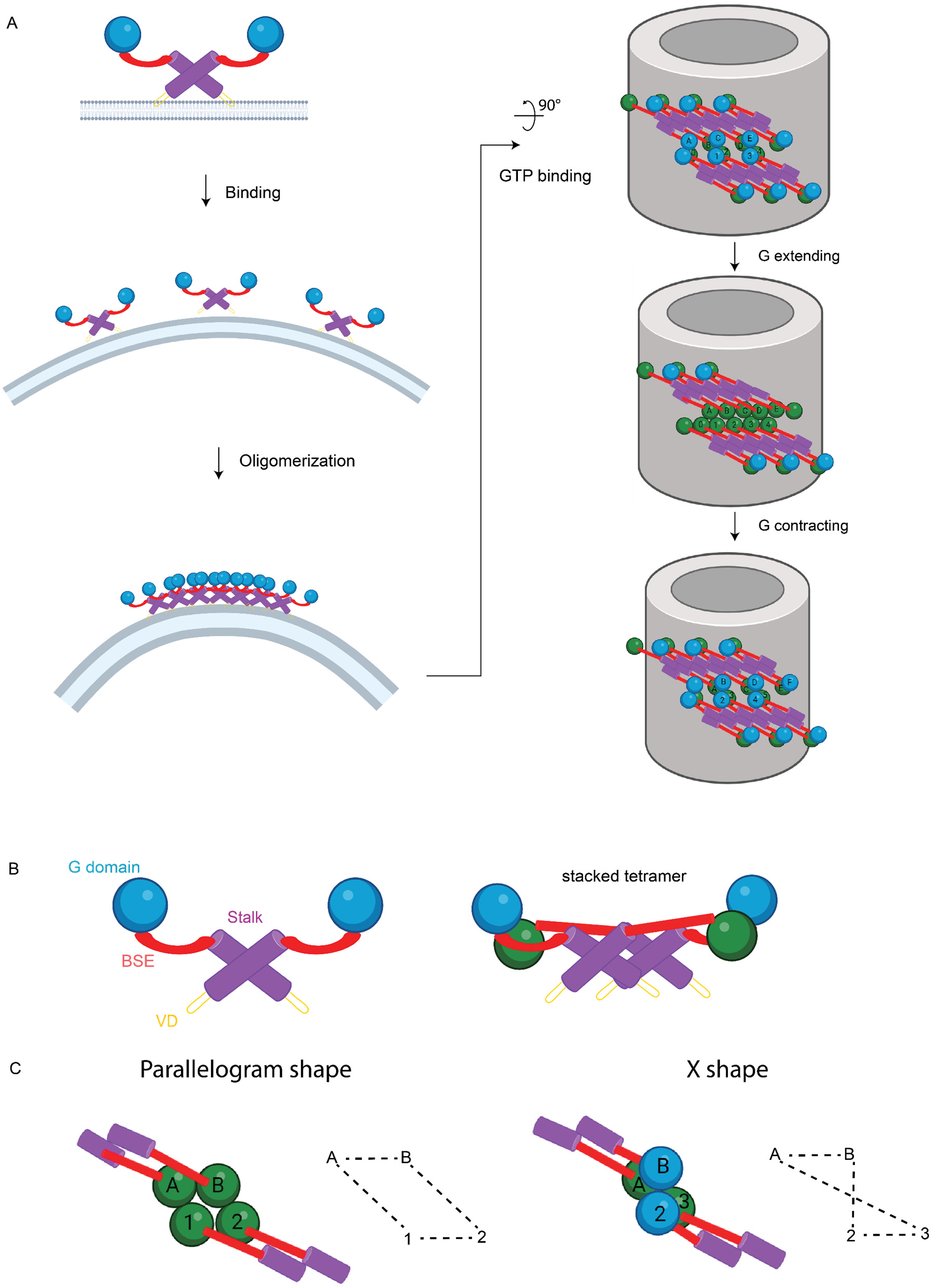
Schematic representation of Drp1 oligomerization on membranes. **(A)** Mitochondria fission model. **(B)** Domain arrangement model of Drp1. **(C)** Connection network between Drp1 rungs. Purple, stalk; Red, BSE; Green, bottom G domain; Blue, top G domain; Brown, VD domain.

## MATERIALS AND METHODS

### Drp1 expression and purification

Drp1 Isoform 3 (UniProt ID O00429-4) was cloned into the pCal-N-EK vector as described previously (Clinton et al. 2016; Clinton, Bauer, and Mears 2020). Drp1-3 was expressed in BL21(DE3) Star *Escherichia coli*. Cells were grown in LB containing 100 µg/mL ampicillin at 18 °C with shaking at 200 rpm for 24 hours after induction with 1 mM isopropyl-1-thio-b-D-galactopyranoside (IPTG). Then, cells were harvested via centrifugation at 4,300 × *g* for 20 minutes at 4 °C. The resulting pellet was resuspended in CalA Buffer (0.5 M L-Arginine pH 7.4, 0.3 M NaCl, 5 mM MgCl_2_, 2 mM CaCl_2_, 1 mM imidazole, 10 mM β-mercaptoethanol) with 1 mM Pefabloc-SC and 100 mg/mL lysozyme. Cells were lysed by sonication on ice. Next, the cell debris was pelleted via centrifugation at 150,000 × g for 1 hour at 4 °C. First, the CBP-tagged Drp1 was purified by affinity chromatography using calmodulin agarose resin (Agilent) that had been pre-equilibrated with CalA Buffer. After the supernatant was loaded onto the column, the resin was washed with 25 column volumes of CalA Buffer. Next, fractions of eluent were collected using 0.5 column volumes of CalB Buffer (0.5 M L-Arginine pH 7.4, 0.3 M NaCl, 2.5 mM EGTA, 10 mM β-mercaptoethanol). Protein-containing fractions were pooled and incubated with GST-tagged PreScission Protease (HRV-3C) overnight at 4 °C to remove the CBP-tag. This solution was concentrated using a 30,000 molecular weight cut-off centrifugal filter (Amicon). This concentrated pool of Drp1 was further purified by size exclusion chromatography (SEC) with an ÄKTA Purifier FPLC (GE Healthcare) and a HiLoad 16/600 Superdex 200 Prep Grade column that had been pre-equilibrated with SEC Buffer (25 mM HEPES (KOH) pH 7.5, 0.15 M KCl, 5 mM MgCl_2_, 10 mM β-mercaptoethanol). All elution fractions containing Drp1 were pooled and concentrated once again, and glycerol (5% final) was added. The purified Drp1 was aliquoted, flash frozen in liquid nitrogen, and stored at −80 °C until use.

### Generation of Drp1 coated nanotubes

All lipid nanotubes utilized here, unless specified otherwise, were comprised of 40% D-galactosyl-beta-1’-N-nervonyl-erythro-sphingosine (GC), 35% phosphatidylethanolamine (PE), and 25% phosphatidic acid (PA). All lipids were purchased from Avanti Polar Lipids, Inc. (Alabaster, AL). Lipids were added to a glass test tube and slowly dried to a thin film using nitrogen gas. The lipid film was then stored in a desiccator for at least one hour to ensure any trace solvent remaining was removed. Then the lipid film was rehydrated with a buffer (200 μL) containing 50 mM HEPES (KOH) pH 7.5 and 0.15 M KCl and heated in 37 °C water bath for ~40 minutes with gentle vortexing every 10 minutes. With these volumes, the final lipid nanotube concentration was 2 mM. The lipid film was placed in a water bath sonicator for 30 seconds and the resulting nanotubes were stored on ice until use. Protein was diluted to 5 μM and incubated at room temperature with lipid nanotubes (500 μM) for at least 30 minutes before adding GMPPCP and 2 mM MgCl_2_.

### Freezing

The sample was frozen using a FEI Vitrobot Mark III. The sample was applied to Holey carbon grids (3.5/1, 400 mesh, Quantifoil) at 4 °C, 100% humidity and incubated on the grid for 60 s. A blot force of 10 and a blot time of 10-12 s was used before plunge freezing into liquid ethane. Samples were stored in liquid nitrogen temperatures before imaging.

### Cryo-EM data collection

Drp1 data was collected in two sessions. For the first dataset where Drp1 was incubated with GalCer tubes at 1 mM GMPPCP, 1,939 images were collected on a Titan Krios (ThermoFisher Scientific). Images were acquired by a DE-64 detector in integrating mode. For the second session where Drp1 is incubated with GalCer tubes at 2 mM GMPPCP, 1,648 images were collected on the same microscope but with the next generation camera, DE Apollo direct detector (Peng et al. 2023). For both datasets, the microscope was operated at 300 kV with a 50 μm C2 aperture and 100 μm objective aperture. The electron beam in nanoprobe mode was aligned through Direct Alignment in the Titan GUI. Coma-free alignment was performed by acquiring a Zemlin tableau in Leginon (Suloway et al. 2005). Cryo-EM images were acquired using the Leginon software and pre-processed using the Appion package (Lander et al. 2009). Apollo camera was set in 8k × 8k super-resolution mode. Movies were collected at a magnification of 47000× with the pixel size of 0.93 Å. Movie frame rate was fixed at 60 frames per second. Movies were collected with a random defocus range of −0.5 μm to −2.0 μm at a detected dose rate of 30 e−/pixel/second (eps), resulting in a total exposure of around 60 e− / Å2.

### Cryo-EM data processing

For the first dataset containing 1 mM GMPPCP, movies were aligned with MotionCor2 (Zheng et al., 2017). The CTF was estimated using CTFFIND4 (Rohou and Grigorieff, 2015) in RELION 3.0. All particles were picked manually, assuring there were no false positives in the dataset. Particles were then sorted into bins by diameter using SPIDER (Frank et al., 1981) scripts and a 2D reference stack. The reference stack was generated by starting with a 2D class average from the particle stack, then pixels were added or removed from the center of the nanotube to respectively increase or decrease the position of the protein decoration to generate 20 bins spanning 43-62 nm in diameter. 15,638 particles from the most populated bins were grouped together for further processing.

For the second dataset (2 mM GMPPCP), movies were aligned MotionCor2 (Zheng et al. 2017). CTF was estimated using CTFFIND4 (Rohou and Grigorieff 2015) supplemented in cryoSPRAC (Punjani et al. 2017). 1,935 micrographs with CTF fit resolution higher than 8 Å were selected for further processing. From the micrographs, 200 filaments particles were picked manually. They were extracted in a small box size of 686. 2D classification is done with a resolution limit of 15 Å, yielding one class with clear GalCer tube features. This 2D class was selected and used as template for a template-based filament tracer in cryoSPARC on 500 micrographs. 98,140 particles were picked in this manner. Another round of 2D classification with resolution limit of 10 Å, yielded 6 good classes with visible tube features. Another round of template-based filament tracer on all micrographs yielded 143,689 particles (box size 960). They are exported using csparc2star.py in pyem package (https://github.com/asarnow/pyem). Particles were binned by 2 for further processing.

### RASTR reconstruction

Particles from both datasets were processed in the same manner unless otherwise specified. Briefly, *psi* angles and shifts were determined using an in-house script first. Then diameters were measured by detecting the membrane signal peaks. Particles of the major peak in diameter distribution histogram were grouped. Random 90 or 270 degrees were assigned as *theta* angles. Random 0 to 360 degrees were assigned as *phi* angles. Particles were reconstructed without alignment using RELION 3.0 (Zivanov et al. 2018) to create an initial average map. The initial map was averaged along the z-axis to further remove features and yielded the final azimuthal average map.

To decide the ROI size, a series of sphere masks with different diameters, all located at the center of GalCer tubes, were created. Separately, ROI was masked out from the azimuthal average map using the sphere mask. Projections were acquired from the masked map using relion_project command in RELION 3.0. These projections were subtracted from raw images using RASTR script. 2D classification was performed to determine the best ROI diameter. Then 4 masks with that diameter positioned at random rotational position around the tube membrane were used to create RASTR particles, resulting in 3 times more particles. These particles were treated as single particles for refinement. 3D classifications were performed, using a masked azimuthal average map as initial model, with C1 symmetry and 8 classes using RELION 3.0. After 25 iterations, good classes with good features were picked for further processing.

For the 1 mM GMPPCP dataset, one good class appeared after 25 iterations. 13,215 particles contributing to the best map from 3D classification were further refined in *cis*TEM with a particle-shaped mask. The Euler angle search was constrained to search only *theta* and *phi*, which converged after four iterations. Then all three Euler angles were refined to convergence using local refinement with gradual increases in the search resolution limit in accordance with the FSC. Local refinement was performed with gradually increased resolution limit, starting at 20 Å using 20% particles, then increasing to 15 Å using 50% particles; each went for 5 iterations. A final 10 iterations of local refinement with resolution limit at 10 Å using all particles were performed to get the final Drp1 map. The final map was post-processed in *cis*TEM and low-pass filtered to remove noise.

For 2 mM GMPPCP dataset, using the featureless azimuthal average map as an initial model, none of the 8 classes yielded a map with clear lattice features. The final map obtained from 1 mM GMPPCP dataset was used as initial model, and a non-uniform refinement was performed. The output map (relaxed state map) represented good stalk features at the center rung but smeared out densities away from center. To further investigate the whole dataset, particles were first reimported into RELION and classified in 3D using the relaxed state map as reference. Three classes appeared with lower resolutions, but consistent stalk features were observed. Differences appeared in the distance of rung spacing were evident. Further refinement could not improve the resolution or quality of the map. Particles were then imported into cryoDRGN 2.2.0. Particles were first binned to box size 120 for initial training, then binned to box size 240 for more accurate training.

Since the edge view of Drp1 tube particles have evident features to distinguish stacked or relaxed state, we used RASTR particles focused on edge to quantify the number of stacked state particles. 90 and 270 degrees were assigned as *phi* angles instead of random 0 to 360 degrees. Subtraction and masking were done in the same manner as described above. Particles were 2D classified in cryoSPARC. Classes with stacked state feature (top G domain) were summed together.

### Model building

The starting tetramer model was built using two different dimer models. For the upper dimer, two monomers the crystal structure were aligned to the G domain interface of a GTPase-GED fusion dimer of Dynamin 1 complexed with GMPPCP (PDB ID: 3ZYC). The lower dimer was built using two monomers from the cryo-EM filament structure with an extended BSE (PDB ID: 5WP9), the G domains were aligned in the same manner described above. The VD was modeled using the Drp1 structure predicted by AlphaFold (Jumper and Hassabis 2022; Jumper et al. 2021). Maintaining the G-G interface, each dimer was fit within the cryo-EM density using rigid body docking in Chimera 1.13 (Pettersen et al. 2004). Four additional chains were docked within the structure to constrain the starting tetramer. In VMD, an all-atom model was generated using the Automatic PSF Builder in VMD28. Files were prepared for NAMD processing following previously described methods (Trabuco et al. 2008). The density map was converted to an MDFF potential. Secondary restraints were applied using NAMD’s extrabonds feature and plugins to preserve secondary structure and to limit artifacts to chiral centers and cis peptide bonds. For simulations, the GScale was set to 0.3, the temperature was set to 300, and the number of timesteps (numsteps) was 25,000. A minimization step was run with a GScale was set to 10, temperature was set to 300 and minimize steps (minsteps) was set to 2000. The model was then minimized and refined using Phenix Real-space refinement (Afonine et al. 2018; Liebschner et al. 2019) to fix clashes and outliers.

### Sub-particle reconstruction

Coordinates of the alpha carbon in amino acid P499 (the starting of VD) were used to calculate the offset for sub-particle centers. Nine offsets were calculated focusing on the center of the stacked state lattice. Nine sub-particles were extracted from every raw particle with box size 128. To create an initial model, *phi* angle offsets were calculated based on the offsets and added/subtracted directly into metadata. A simple relion_reconstruct yielded the initial map. Sub-particles were refined without imposing symmetry in *cis*TEM suite.

### Tomogram Reconstruction and Segmentation

Tilt stacks were processed in RELION 5.0 (Burt et al. 2024) using the IMOD wrapper to apply a patch-tracking. The IMOD results were used to generate reconstructions with a SIRT-like filter to enhance contrast in the IMOD 4.12.53 (Mastronarde 1997) etomo application. Reconstructions were imported into Dragonfly 2024.1 (Dragonfly 2024.1 non-commercial software. Comet Technologies Canada Inc., Montreal, Canada; software available at https://dragonfly.comet.tech/). Three imaging filters were applied to further enhance contrast: a histogram equalization, a gaussian high pass smoothing filter, and a sharping filter. Decorated tubes were manually segmented and visualized in ChimeraX.

### Malachite Green Colorimetric Assay

The basal GTPase activity of Drp1 was measured using a colorimetric assay to detect released phosphate, as described previously (Clinton, Bauer, and Mears 2020; Clinton et al. 2016). Briefly, Drp1 (500 nM final) was diluted to 2.4X with Assembly Buffer (25 mM HEPES (KOH) pH 7.5, 150 mM KCl, 10 mM β-mercaptoethanol). To start the reaction, 3X GTP/MgCl_2_ (1 mM and 2 mM final, respectively) was added to the Drp1 incubated with either 4X lipid (150 μM final) to calculate the lipid-stimulated rates or only Assembly Buffer to calculate the rate for the protein alone in solution. The reaction was carried out at 37°C. At the chosen time points, a sample aliquot was taken and quickly added to EDTA (100 mM final) to stop the reaction. The time points used to calculate the rate for the protein alone in solution were 0, 5, 10, 15, 45, and 60 min. For the lipid-stimulated rates the time points were 0, 2, 4, 6, 8, and 10 min. After collecting all time points, Malachite Green Reagent (1 mM malachite green carbinol, 10 mM ammonium molybdate tetrahydrate, 1 N HCl) was added to each sample, and the absorbance at 650 nm was measured using a VersaMax microplate reader (Molecular Devices).

### Sedimentation Assay

To quantify Drp1 oligomerization, a sedimentation assay was conducted similar to what has been described previously (Francy et al. 2015; Mears and Hinshaw 2008). Large oligomers formed by Drp1 samples, in the presence of ligands, were found in the pellet after a medium speed centrifugation. Specifically, protein was diluted in HEPES KCl buffer to 2 μM, and specified WT and mutant samples were incubated at room temperature with cardiolipin-containing lipid nanotubes or liposomes (300 μM) for at least 60 minutes. The mixtures were then spun at 15,000 rpm for 10 min in a tabletop centrifuge (Eppendorf). The supernatant and pellet fractions were separated, collected, and immediately mixed with SDS-PAGE loading dye (Bio-Rad) and heated briefly at 100 °C. These samples were run on an SDS-PAGE gel and stained with an InstantBlue Coomassie dye (Expedeon). Gels were scanned using an Odyssey XF Imaging System (Li-Cor) and densitometry analysis was done using ImageJ.

## ACKNOWLEDGEMENTS

K.R. discloses support for the research of this work from the National Institute of General Medical Sciences [F31 GM139324]. S.M.S. discloses support for the research and publication of this work from National Institute of General Medical Sciences [R01 GM108753] and [R01 GM143805]. J.A.M. discloses support for the research and publication of this work from National Institute of General Medical Sciences [R01 GM125844].

Molecular graphics images were produced using the UCSF Chimera package from the Resource for Biocomputing, Visualization, and Informatics at the University of California, San Francisco (supported by NIH P41 RR-01081).

## AUTHOR CONTRIBUTIONS

Conceptualization: Peng, R., Rochon, K., Stagg, S.M., and Mears, J.A.; Funding acquisition: Rochon, K., Stagg, S.M., and Mears, J.A.; Sample Preparation: Rochon, K. and Mears, J.A.; Imaging and Data Acquisition: Peng, R., Stagg, S.M., and Mears, J.A.; Image Processing Methodology: Peng, R. and Stagg, S.M.; Model building: Rochon, K.; Supervision: Stagg, S.M., and Mears, J.A.; Writing, reviewing and editing: Peng, R., Rochon, K., Stagg, S.M., and Mears, J.A.

## Notes

### Competing Interest Statement

The authors have declared no competing interest.

### Summary of Updates

Introduction changed to remove a paragraph. New reference added.

