## Supplemental figures for "The Structure of the Drp1 Lattice on Membrane"

### Supplementary Information

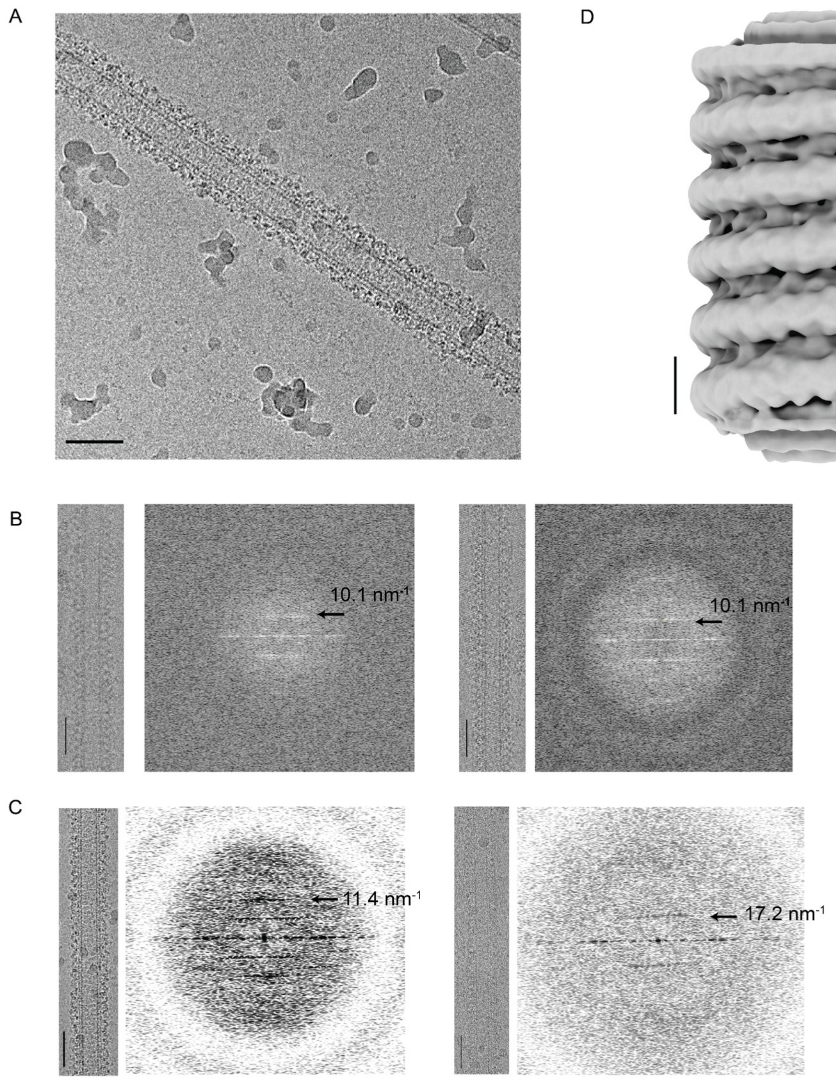

**Figure S1. Drp1 helical reconstruction. (A)** Representative micrograph of Drp1 decorated GalCer tube. **(B)** Two representative segments of Drp1 incubated with GalCer at 1mM GMPPCP. Left, segment image. Right, zoomed Fourier transform of the segment. **(C)** Two representative segments of Drp1 incubated with GalCer at 2mM GMPPCP. Left, segment image. Right, zoomed Fourier transform of the segment. **(D)** Helical reconstruction using helical symmetry parameters estimated in Fig. 1B. Scale bar, 50nm for A, B, C. Scale bar, 20nm for D.

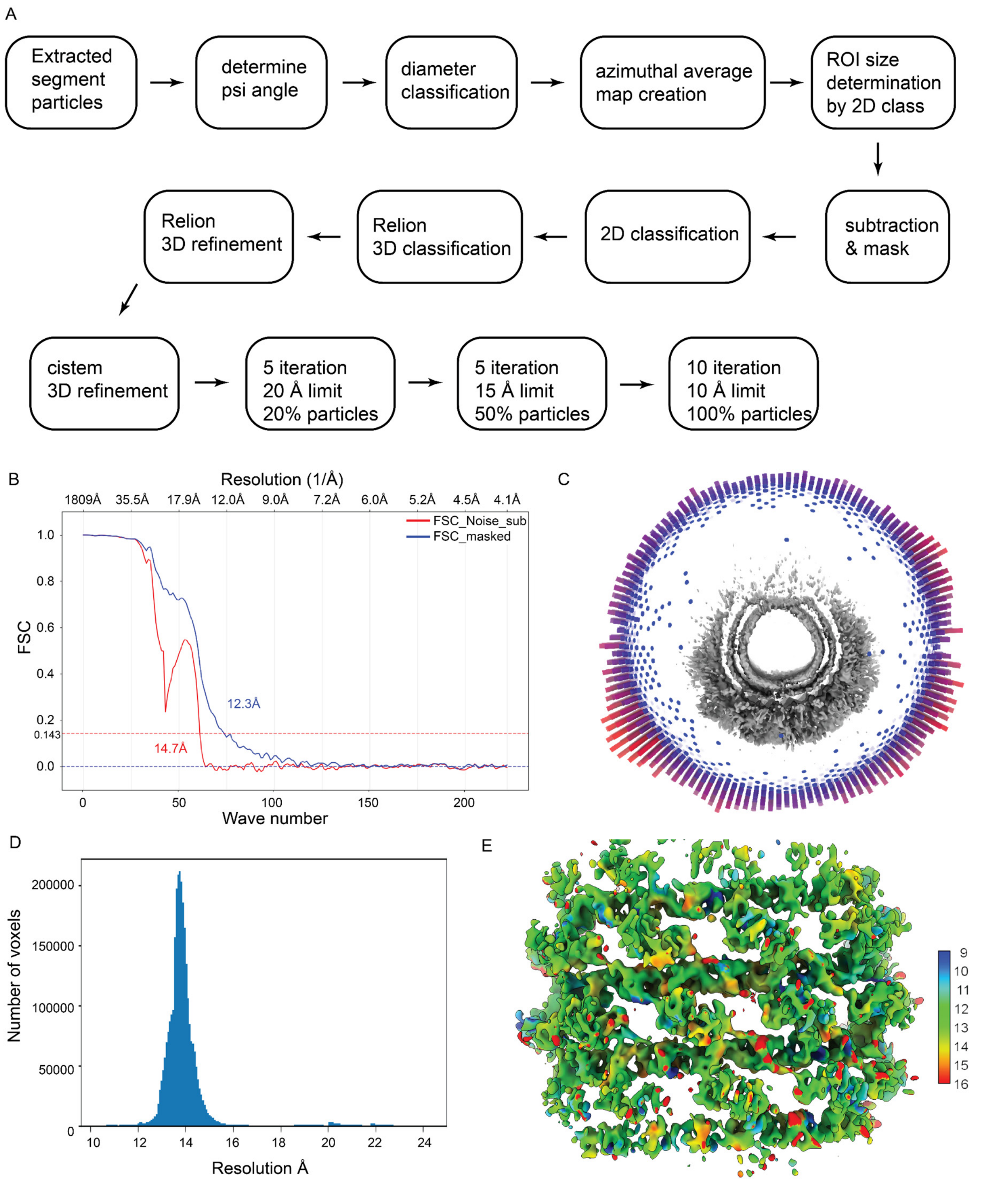

**Figure S2. Data processing of stacked state Drp1 lattice**. **(A)** Workflow of data processing. **(B)** FSC curve of stacked state Drp1 lattice reconstruction. Red, phase randomized FSC. Blue, FSC with mask of protein lattice shape. **(C)** Angle distribution of stacked state Drp1 lattice reconstruction. **(D)** Local resolution counts of stacked state Drp1 map. **(E)** Stacked state map colored by local resolution index.

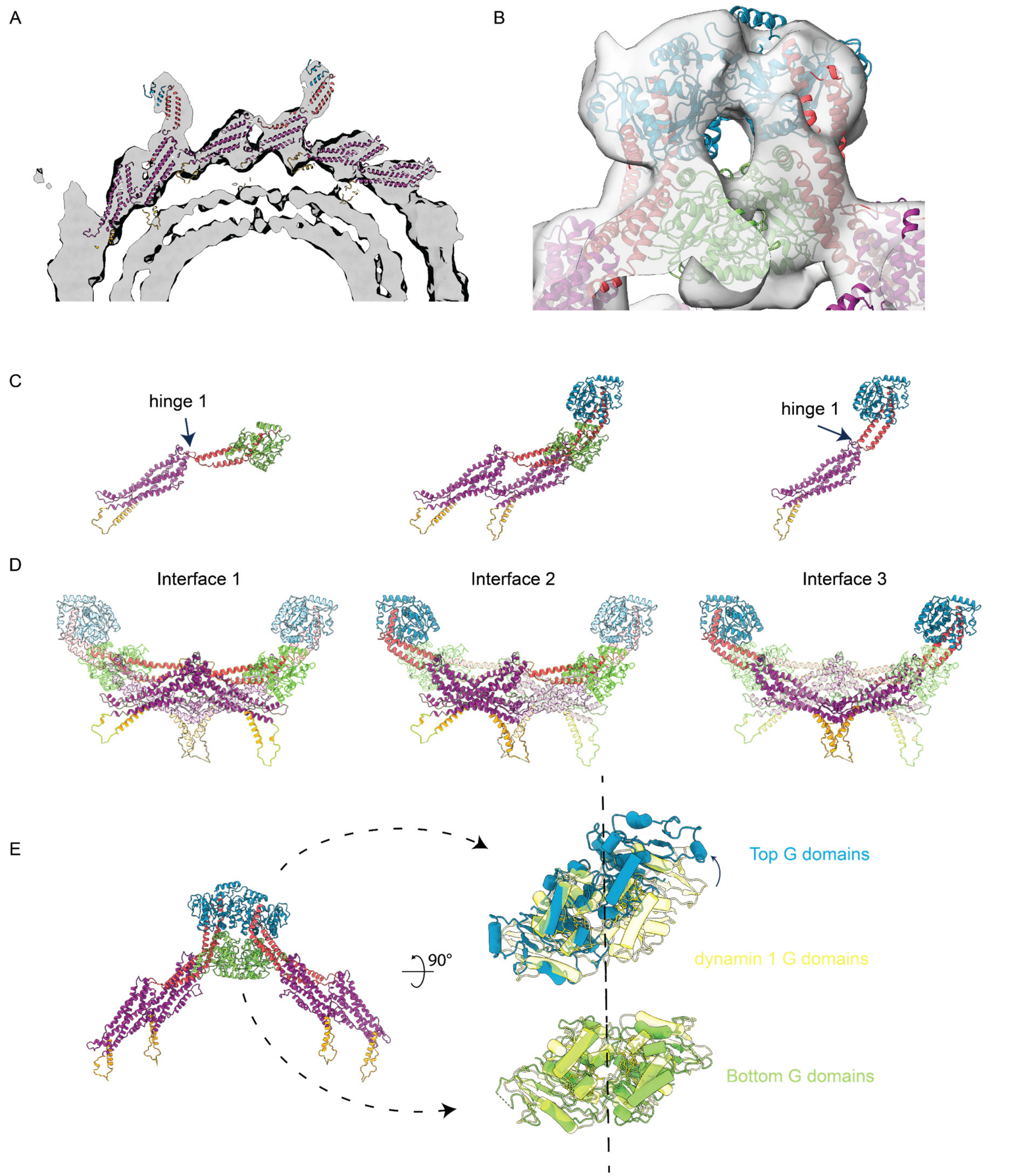

**Figure S3. Model details of stacked state Drp1 lattice. (A)** Stalk domain arrangement. A slice of stacked state Drp1 cryo-EM map focusing on one side of the stalk domain filament. **(B)** G domain fitting. **(C)** Poses between stalk domain and BSE. Green G domain represent extended conformation. Blue G domain represent contracted conformation. **(D)** Interfaces of stalk domains. Left, interface 1. Middle, interface 2. Right, interface 3. **(E)** GG interface of stacked state Drp1 lattice. Right images are taken as looking from outer radial position to the center of tubules. Transparent yellow represent GG domains in dynamin 1 (PDB ID: 6DLU). Left G domains are fitted together.

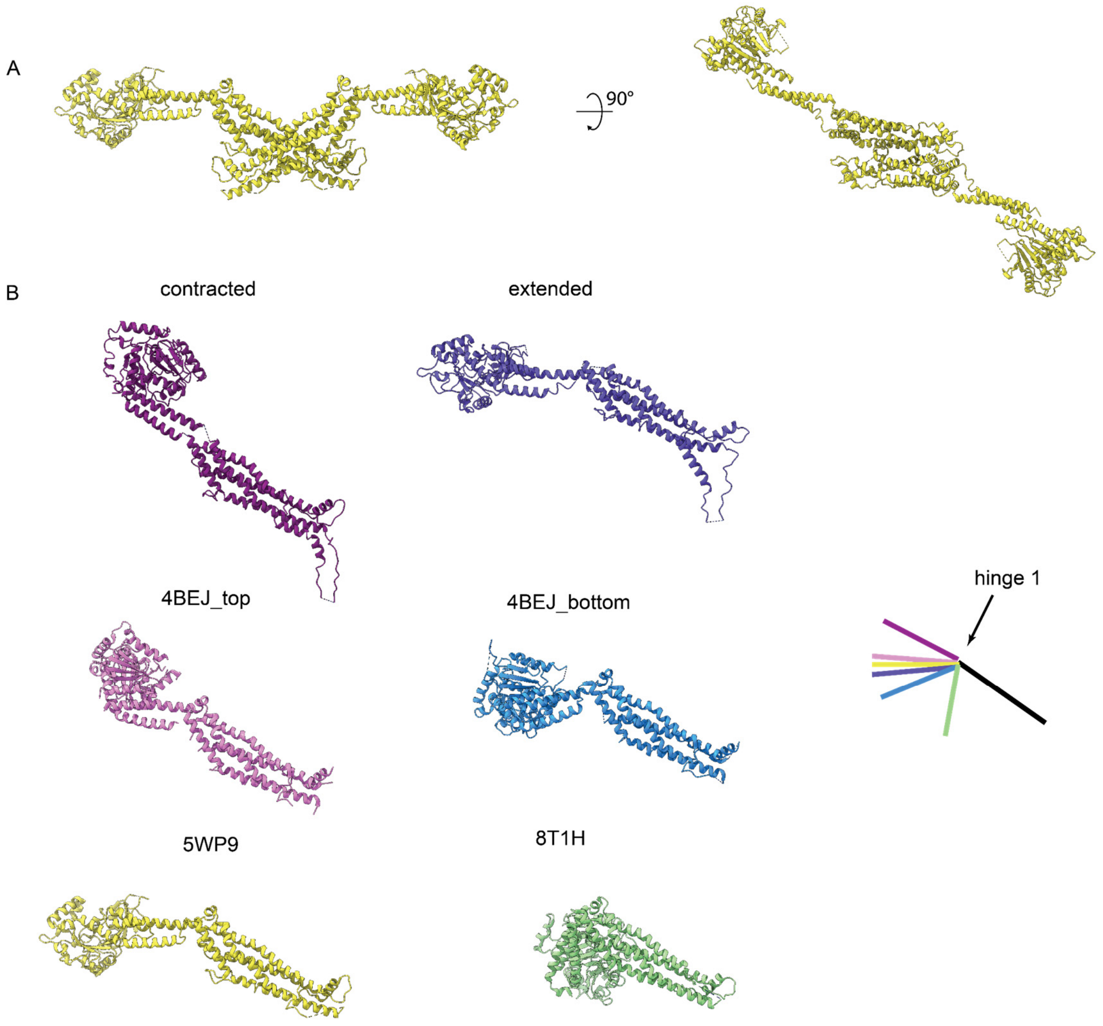

**Figure S4. Hinge 1 conformation comparison. (A)** 5WP9 dimer showing orientations taken for comparison. **(B)** Individual monomers comparison. All models are superimposed on stalk region.

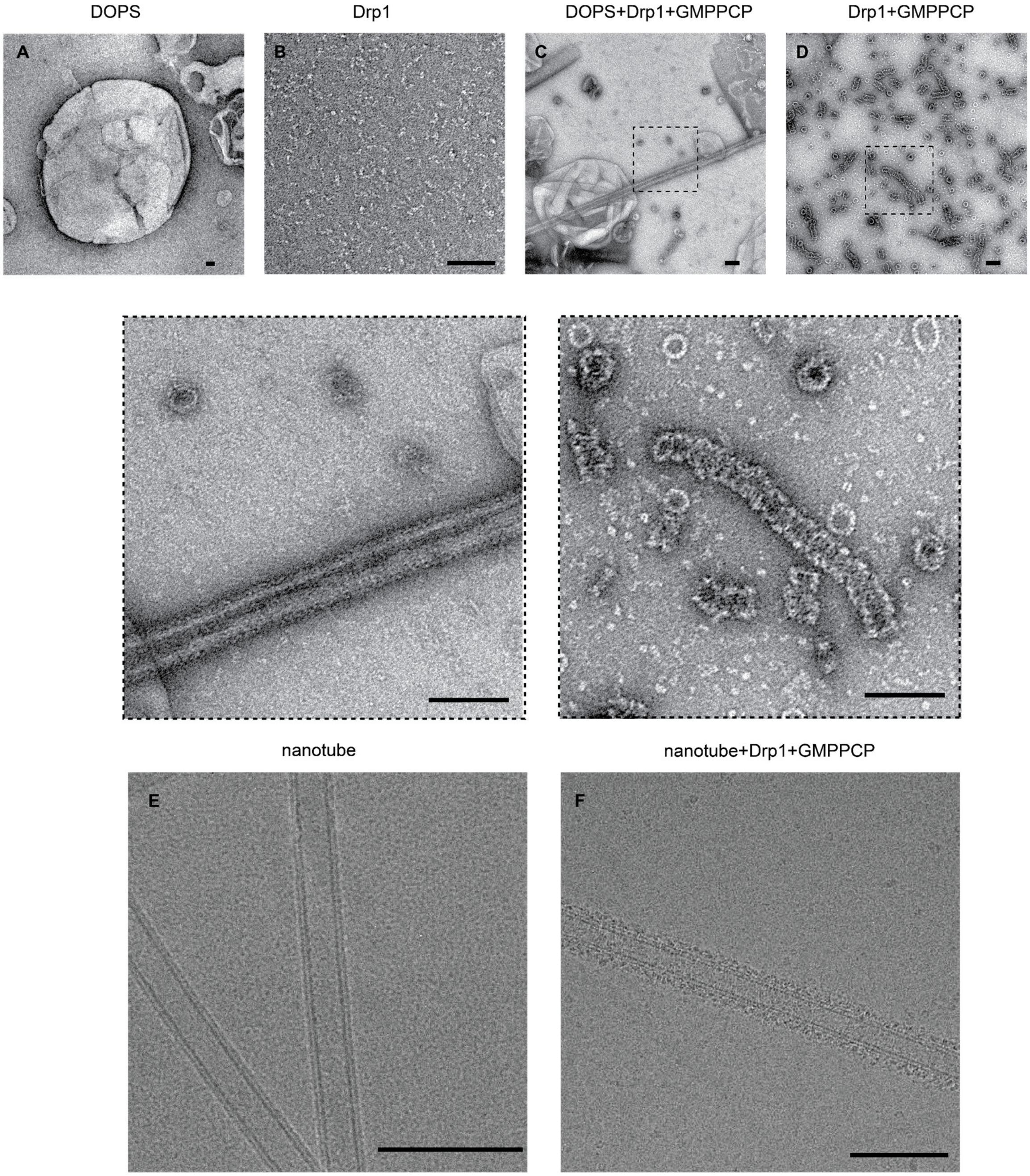

**Figure S5. Drp1 assembly on membrane.** Negative staining image of DOPS liposome **(A)**, Drp1 alone **(B)** DOPS incubated with Drp1 in the presence of 1mM GMPPCP **(C)** Drp1 incubated with 1mM GMPCP **(D).** (**E)** Cryo-EM image of naked GalCer tube. (40% GalCer, 35% PE, 25% PA)**. (F)** Cryo-EM image of Drp1 decorated GalCer tube. Scale bar, 100 nm.

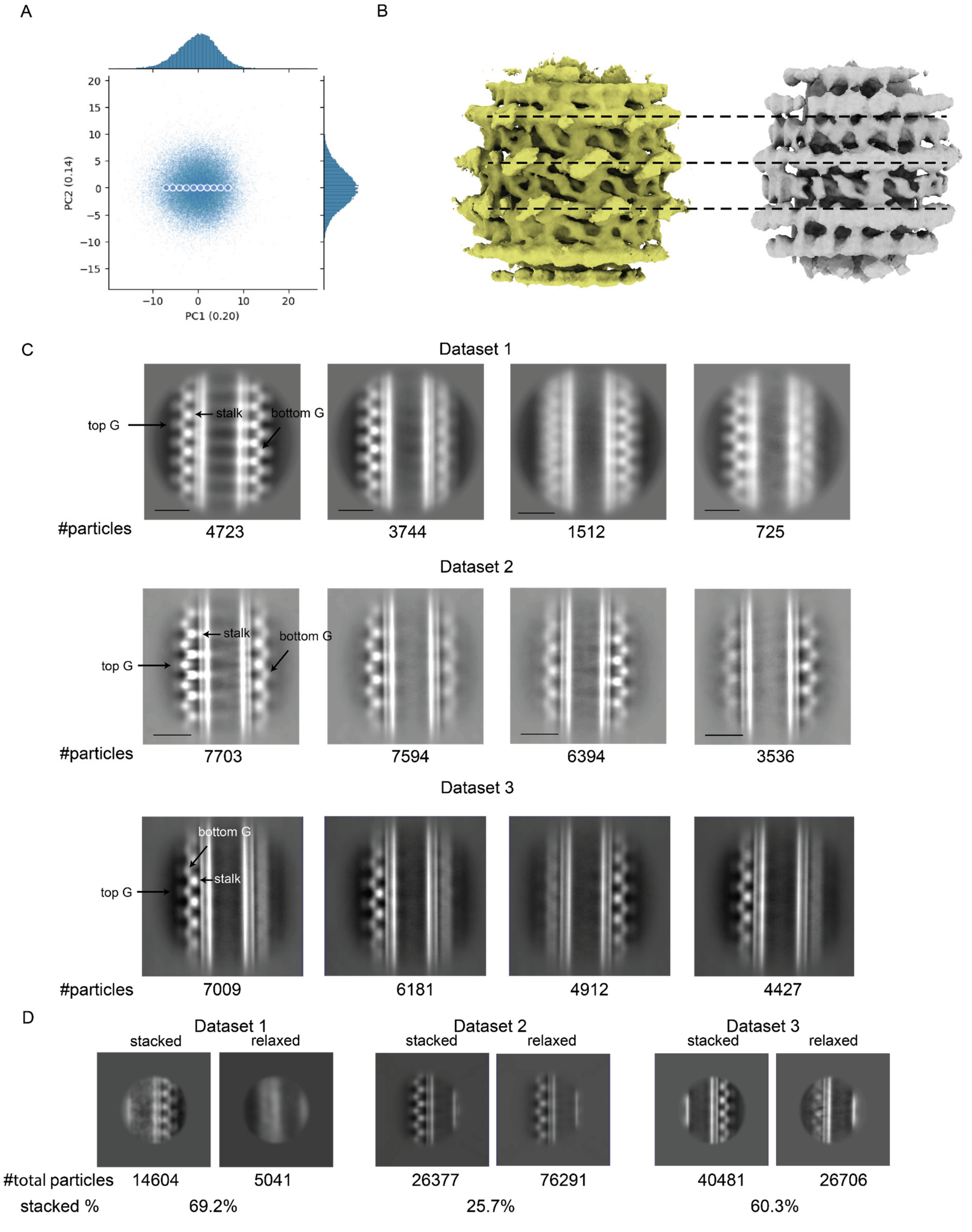

**Figure S6. Heterogeneity in Drp1 datasets. (A)** Latent space representation of dataset 2 particles using cryoDRGN. Points indicates maps used in Supplementary Video 2. **(B)** Rung spacing comparision. **(C)** Representative 2D class averages of raw filament particles. Scale bar 20nm. **(D)** Representative 2D class averages of RASTR particles focused on edge views. Particle numbers from 2D classes with specified features are summed together. Percentage of particles with stacked top G domains are listed below.

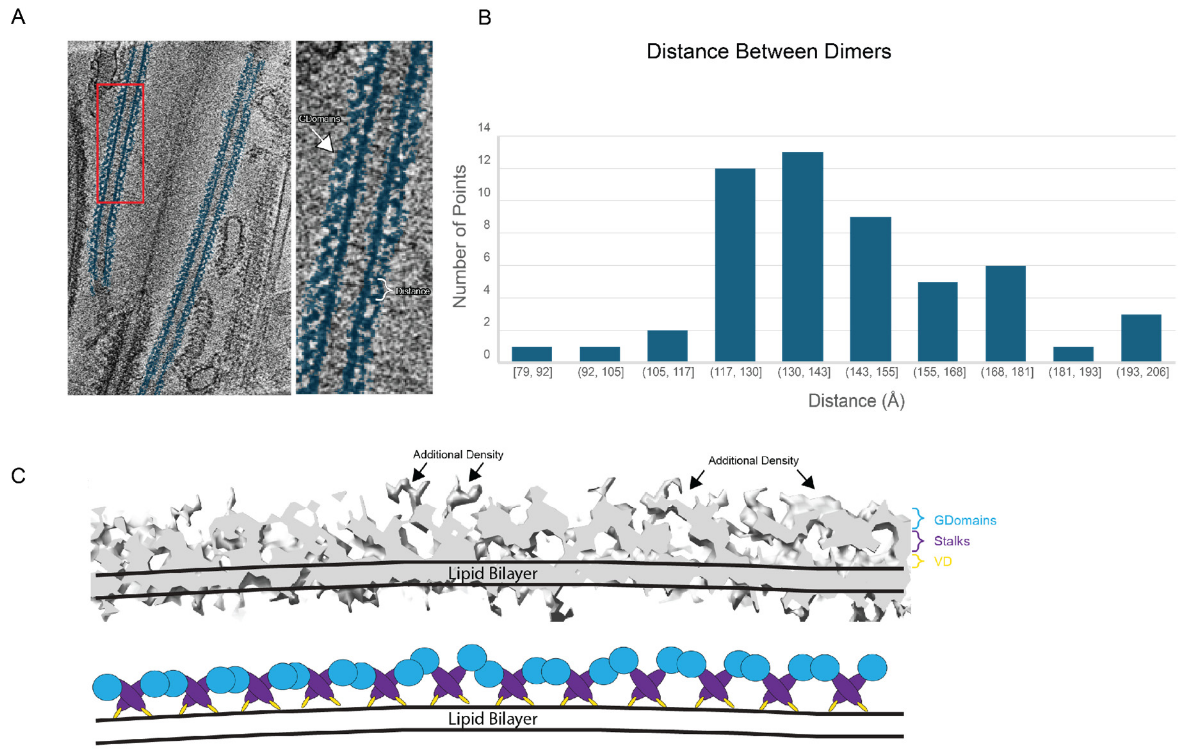

**Figure S7. Cryo-ET on Drp1 decorated GalCer tubes. (A)** Tomography of Drp1 decorated GalCer. Protein features colored blue. **(B)** Distance quantification of tomography reconstruction. Distance is defined as the spacing between adjacent stalks. **(C)** Tomography map and schematic models.

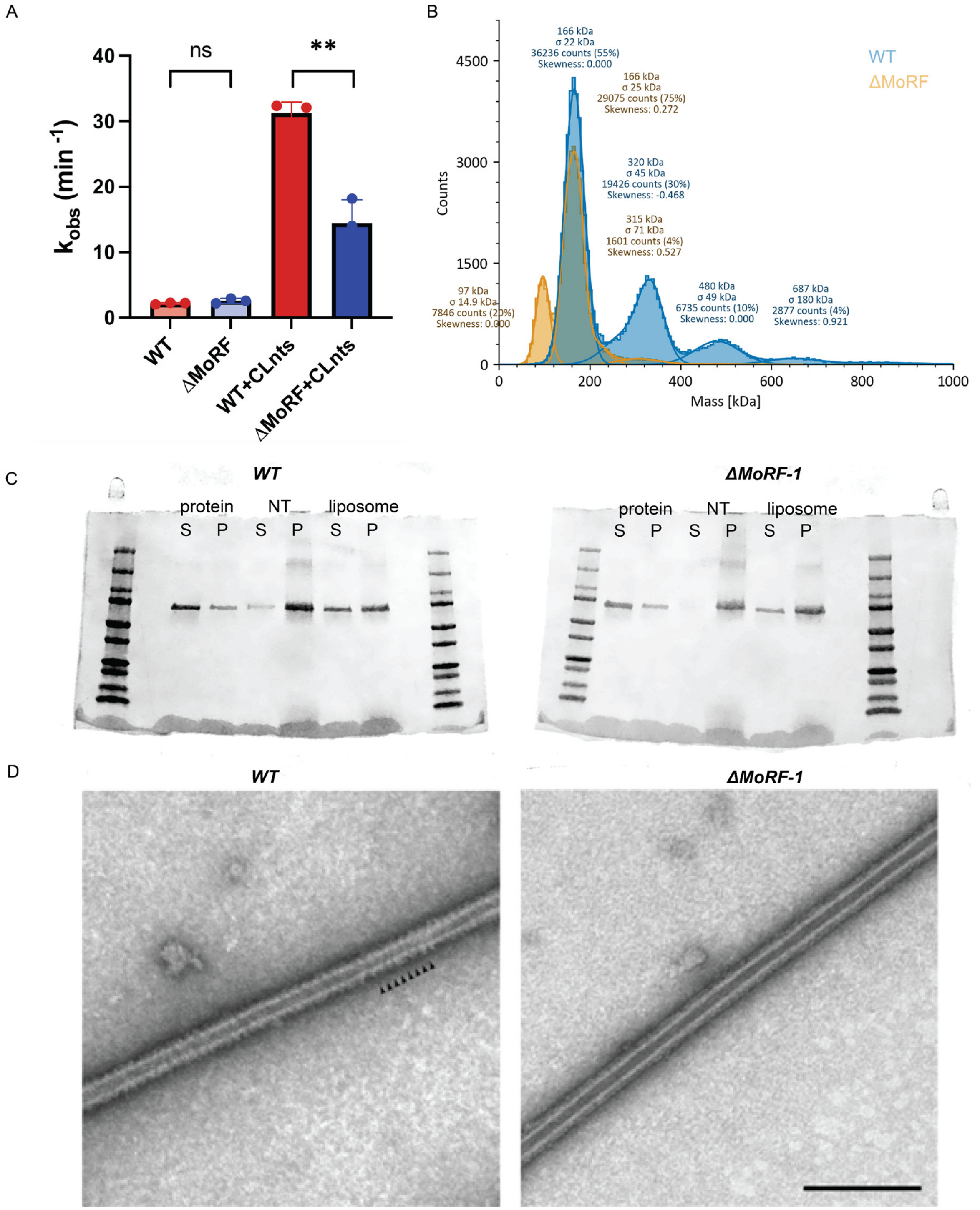

**Figure S8. MoRF-1 deletion. (A)** GTPase activity upon membrane stimulation. **(B)** Molecular weight determined by mass photometry. **(C)** Liposome co-sedimentation assay. S for supernatant. P for pellet. **(D)** Negative staining images of Drp1 decorated membrane tubules. Small arrowheads highlight regular striations in the lattice. Scale bar, 200 nm.

Supplementary Table 1. Cryo-EM data collection, refinement, and validation statistics

| | Drp1 Asymmetric Tetramer (EMDB-43045) (PDB 8V8T) |
|---|---|
| **Data collection and processing** | |
| Voltage (kV) | 300 |
| Electron exposure (e–/Å$^2$) | 58.49 |
| Defocus range (μm) | -0.8 to -2.0 |
| Pixel size (Å) | 2.02 |
| Symmetry imposed | C1 |
| Initial particle images (no.) | 51,102 |
| Final particle images (no.) | 13,215 |
| Map resolution (Å) | 11.37 |
| FSC threshold | 0.143 |
| **Refinement** | |
| Initial model used (PDB code) | 4BEJ, 5WP9 |
| Model composition | |
| Non-hydrogen atoms | 22,382 |
| Protein residues | 2918 |
| Ligands | 0 |
| R.m.s. deviations | |
| Bond lengths (Å) | 0.003 |
| Bond angles (°) | 0.706 |
| Validation | |
| MolProbity score | 2.81 |
| Clashscore | 12.51 |
| Poor rotamers (%) | |
| Ramachandran plot | |
| Favored (%) | 89.09 |
| Allowed (%) | 9.64 |
| Disallowed (%) | 1.27 |
